# HPL-2/HP1 and MET-2/SETDB1 bind distinct co-factors that promote heterochromatic foci, gene repression and organogenesis independently of H3K9 methylation

**DOI:** 10.1101/2024.09.29.615660

**Authors:** Colin E. Delaney, Lisa Fol, Jia Emil You, Stefan B. Gruenewald, Eugenio Ferrario, Susan M. Gasser, Martin M. Moeckel, Lukas Stelzl, Jan Padeken, Susan E. Mango

## Abstract

In differentiated cells, genome segregation into heterochromatin and euchromatin is mediated by modified histones, which recruit so-called reader proteins. Surprisingly, many histone modifiers remain functional in the absence of catalytic activity, but the underlying mechanism remains unclear. To explore this puzzle, we examined the relationship between *C. elegans* MET-2/SETDB1, a histone H3 lysine (H3K9me) methyltransferase that also has non-catalytic roles, and the canonical H3K9me reader HPL-2 (HP1). We show that HPL-2 represses transcription and supports organogenesis independently of H3K9me binding, whereas complete loss of *met-2* and *hpl-2* causes severe transcriptional and developmental defects. MET-2 and HPL-2 rely on different binding partners - the disordered protein LIN-65/ATF7IP and the multi-zinc finger protein LIN-13, respectively – for localization and function. The results suggest that HPL-2 can operate through alternative protein interactions, and that HPL-2 and MET-2 function in parallel, H3K9me-independent pathways, with H3K9me acting as a reinforcing but non-essential contributor to these processes.

## Introduction

Tissue differentiation in multicellular organisms is accompanied by the segregation of the genome into euchromatin and heterochromatin. Open euchromatin is accessible to transcriptional activators while compacted heterochromatin restricts transcription and relocates to the nuclear periphery and nucleolus (1). Distinct sets of post-translational modifications of histones (PTMs) are enriched in these genomic compartments and play important roles in transcriptional regulation. Methylation of histone H3 on lysine 9 (H3K9me) is a hallmark of transcriptionally silenced heterochromatin, marking satellite and simple repeats, transposable elements, and inactive tissue-specific genes (2). Loss of H3K9me-specific lysine methyltransferases (HMTs) that write H3K9me via their SET domains (<u>S</u>uppressor of Variegation, <u>E</u>nhancer of Zeste, <u>T</u>rithorax domain) is associated with defects in tissue differentiation, aging, and altered tumor immunogenicity (3–8). Once H3K9me is deposited, reader proteins like the highly conserved HP1 (heterochromatin protein 1) bind specifically to H3K9me2/me3 via their chromodomain (9, 10). HP1 binding of H3K9me, followed by its dimerization via its chromoshadow domain, is a well-established model for heterochromatin compaction and gene silencing.

While the prevailing view suggests HP1 binds H3K9me to function, recent studies have suggested a more nuanced relationship. HP1 can interact with chromatin through H3K9me binding but also via DNA and RNA binding to the hinge domain (11–14), and single molecule live imaging demonstrates dynamic H3K9me-independent HP1-nucleosome interactions (15). These observations suggest H3K9me independent activities, but to what extent PTM-dependent and -independent mechanisms contribute to HP1 function remains unclear. Whether or how H3K9me-independent interactions are linked to the demonstrated ability of HP1 to phase-separate in vitro, and show condensate-like fusing of foci in vivo, is a subject of active investigation (16–18).

In the nematode *C. elegans*, the relationship between HP1 homolog HPL-2 and H3K9me is similarly ill defined. HPL-2, MET-2/SETDB1, and H3K9me2 map to the same areas of the genome and repress repetitive elements in whole animal ChIP-seq experiments (19). However, overexpressed HPL-2 can bind target loci in embryos lacking all H3K9me, albeit at a weaker level (20). Further, HPL-2 acts redundantly with the SETDB1 homolog MET-2 in germline stability and vulval cell differentiation, suggesting independent functions (21, 22). The histone methyltransferase MET-2 is required for H3K9me-mediated gene silencing (23, 24), but also possesses a catalytic-independent function that maintains hypoacetylation of heterochromatin, mitigates phenotypes associated with the loss of H3K9me, and retains the ability to repress some genes (25). During interphase MET-2 forms dynamic heterochromatic foci that are independent of H3K9me and instead rely on interaction with the largely unstructured protein LIN-65/ATF7IP (26–28). Interestingly, overexpressed HPL-2 has also been reported to localize to sub-nuclear foci (29); however, it is unclear how MET-2/SETDB1, H3K9me deposition, and HPL-2/HP1 functionally interact in transcriptional silencing and tissue differentiation.

Here, we interrogate the relationship between HPL-2/HP1, H3K9me, and MET-2/SETDB1 in localization, gene silencing, protein dynamics, and tissue differentiation. We find that while HPL-2 co-localizes with MET-2 in subnuclear heterochromatic foci, their recruitment to these foci is independent from each other. The independent targeting to heterochromatin foci correlates with a redundant role of MET-2 and HPL-2 in gene repression and restricting cell fate. Strikingly, HPL-2 localization does not require H3K9me binding. Instead, HPL-2 localization depends on physical interaction with the multi-zinc finger protein LIN-13 mediated by the chromoshadow domain. H3K9me binding plays only a minor role in HPL-2 mediated transcriptional repression and is dispensable for cell fate. These findings argue that multivalent protein-protein interactions may constitute a generalizable mechanism for heterochromatin function as an additional layer of regulation to histone PTMs.

## Results

### MET-2 and HPL-2 colocalize and are independently recruited into heterochromatic foci

To begin dissecting the relationship between MET-2/SETDB1, H3K9me, and HPL-2/HP1, we asked if endogenous HPL-2 forms foci and if so, how MET-2 and HPL-2 foci interact by imaging live, prebean-stage *C. elegans* embryos. We monitored HPL-2 protein using a strain expressing HPL-2 CRISPR-tagged with mKate2 and 3xFLAG inserted into the unstructured hinge region (Figure 1A) (30). The *hpl-2* gene encodes 2 isoforms (31). The major isoform accounts for 93% of total protein and exhibits the typical HP1 structure of chromodomain (CD)-hinge-chromoshadow domain (CSD), while the less abundant isoform possesses an abbreviated CSD and an extended C terminus (Fig. 1A). The internal HPL-2::mKate2 fusion tags both isoforms.

**Fig. 1.**
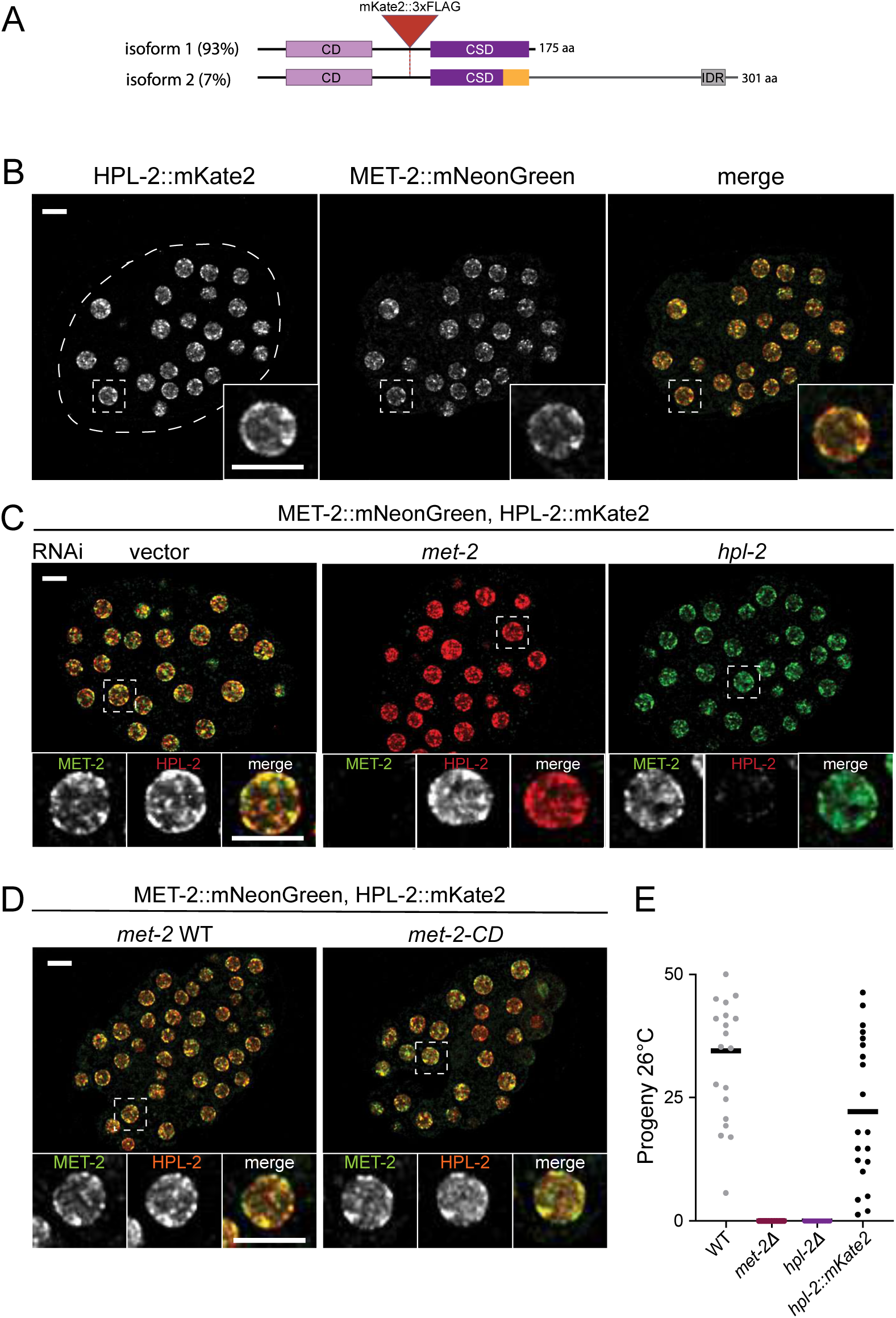
MET-2 and HPL-2 colocalize via independent recruitment into heterochromatic foci. (**A**) Schematic of HPL-2 protein isoforms expressed from the same locus, with isoform percentage of total protein indicated in parentheses as determined by Western blot (mean, N = 3; see also Fig. 3F, S4B). Red triangle = mKate2::3xFLAG insertion site in the hinge domain. (**B**) Representative images of MET-2::mNeonGreen co-expressed with HPL-2::mKate2. Dashed oval indicates the circumference of the embryo. Dashed boxes indicate example nuclei enlarged below. Scale bar, 5 μm. N = 3, n = 30. **(C)** Representative images of embryos co-expressing MET-2::mNeonGreen with HPL-2::mKate2 treated with *met-2*, or *hpl-2* RNAi or empty vector control. Scale bar, 5 μm. N = 3, n = 30. **(D)** Representative images of MET-2::mNeonGreen co-expressed with HPL-2::mKate2 in WT, *met-2-CD*, and *hpl-2-FFAA* backgrounds. Scale bar, 5 μm. N = 3, n = 30. **(E)** Mean progeny per worm at 26°C in WT, *hpl-2Δ*, and *hpl-2::mKate2::3xFLAG*. N = 2, n = 60 egg layers. Bars indicate mean value of each genotype.

HPL-2::mKate2 was constitutively nuclear and concentrated into subnuclear foci (Fig. 1B). We then co-expressed HPL-2::mKate2 with MET-2 endogenously tagged with mNeonGreen (32), and observed that HPL-2 foci to a large extent colocalized with MET-2 foci (Fig. 1B). Knockdown of either *met-2* or *hpl-2* led to the loss of green or red fluorescence, respectively, indicating that the signal was specific (Fig. 1C). Surprisingly, the depletion of *met-2* did not result in a loss of HPL-2 foci (Figure 1C). We previously showed that MET-2 foci form independent of H3K9me using a MET-2 mutant deficient in catalytic activity (*met-2-CD*) (25). Similarly, HPL-2 foci persisted in *met-2-CD* embryos (Fig. 1D). Loss of heterochromatin results in temperature-dependent sterility (19, 24, 33–35). We confirmed that the strain expressing HPL-2::mKate2 remained viable and fertile at 26°C unlike *hpl-2Δ* animals (Fig. 1E), indicating that HPL-2::mKate2 is functional and raising the question how foci may contribute to HPL-2 function.

Factors downstream of MET-2 in the H3K9me pathway, *set-25* and *lin-61* (36), were dispensable for both MET-2 and HPL-2 foci (Fig. S1A). Loss of *hpl-1*, another HP1-like gene, does not exhibit the somatic and germline physiological defects seen in *met-2* or *hpl-2* mutant animals (31), and *hpl-1* was not necessary for MET-2 or HPL-2 foci (Fig. S1A). Further, endogenously tagged HPL-1 was observed to be expressed at the minimum limit of detection in embryos, much lower than HPL-2 levels, prior to upregulation in late embryogenesis (Fig. S1B and data not shown, (30)). Therefore, we focused on HPL-2, finding that HPL-2 and MET-2 form foci independent of each other. This result suggests two parallel pathways for subnuclear localization of heterochromatin factors MET-2 and HPL-2.

### MET-2 and HPL-2 repress transcription via parallel pathways

To begin dissecting how HPL-2 and MET-2’s independent targeting to foci may correlate with their functions, we examined the effect of HPL-2 and MET-2 on transcription in *C. elegans* embryos. The well-characterized repetitive heterochromatic array *gwIs4* expresses nuclear GFP and is repressed by MET-2, SET-25, and HPL-2 (37, 38). Double deletion of both *met-2* and *set-25* eliminates all detectable H3K9me and leads to desilencing of the array (Fig. 2A) (38). Knockdown of *hpl-2* using RNAi led to a significant increase in GFP expression not only in WT, but also in *met-2 set-25* double mutants (Fig. 2A), indicating that HPL-2 has a silencing function independent of H3K9me binding. Consistently, double deletion of *met-2* and *hpl-2* shows synthetic phenotypes, suggesting parallel functions (Fig. 2B and (21, 22)).

**Fig 2.**
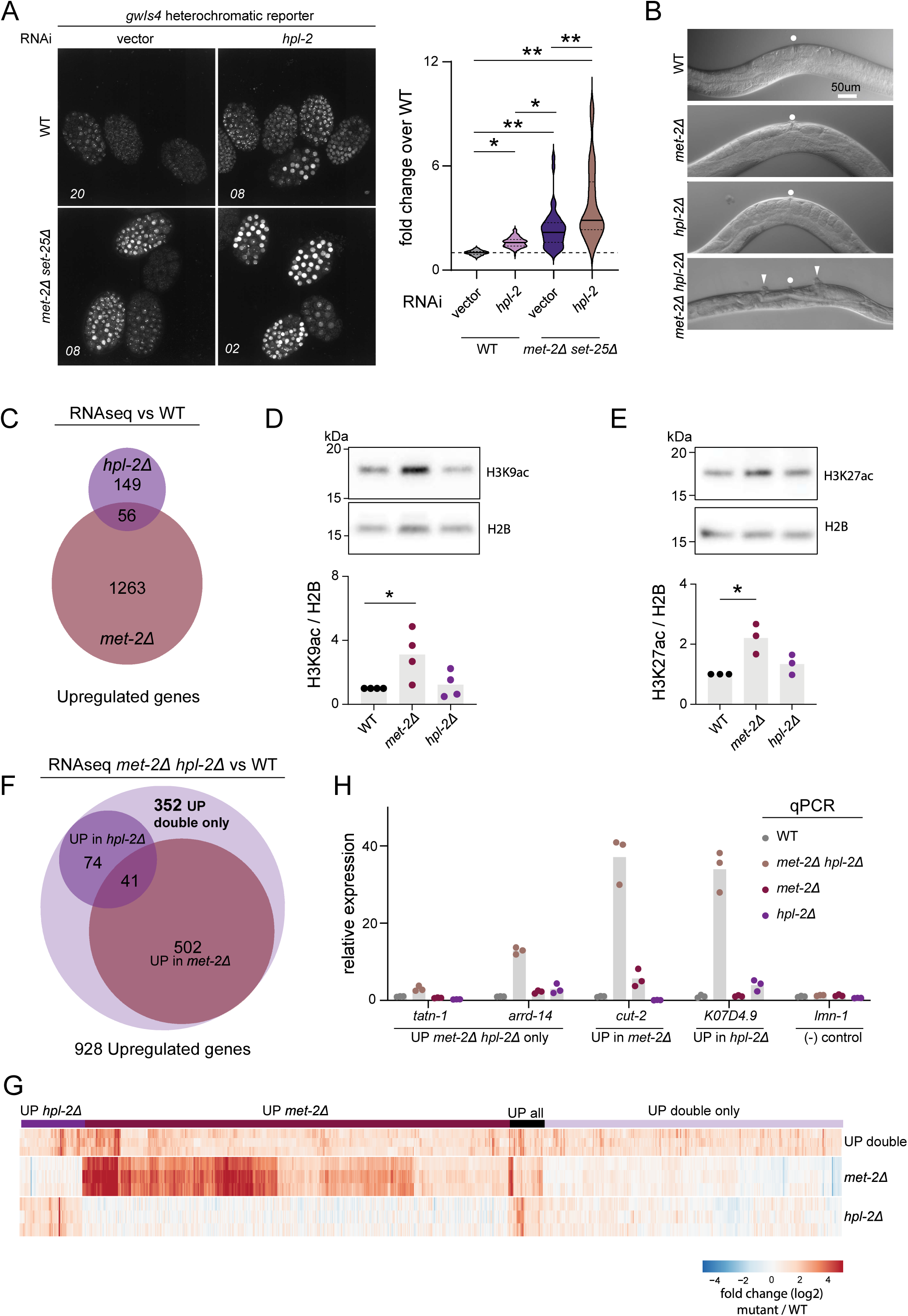
H3K9me and HPL-2 act independently to repress transcription. (**A**) Representative images and quantification of GFP signal from the *gwIs4* heterochromatic array. N = 3, n(embryos): WT vector = 99, WT *hpl-2* RNAi = 134, *met-2Δ set-25Δ* vector = 39, *met-2Δ set-25Δ hpl-2* RNAi = 53. *p < 3x10^-4^, **p < 10^-9^ by one-way ANOVA, followed by Bonferroni post hoc test. (**B**) Representative images of vulva development in adult animals. Normal vulvae = circle, multivulva defects = arrowheads. Quantification reported previously (21). (**C**) Venn diagram showing overlap of *hpl-2Δ* and *met-2Δ*-derepressed genes measured by RNAseq. N = 3, FC > 2, FDR < 0.05. (**D**) Representative western blot and quantification of acetylated H3K9 normalized to histone H2B in WT, *hpl-2Δ* and *met-2Δ* single mutants. *N* = 4. *p = 0.034 by two-sided *t*-test. (**E**) Representative western blot and quantification of H3K27 acetylation normalized to histone H2B in WT, *hpl-2Δ* and *met-2Δ* single mutants. *N* = 3. *p = 0.014 by two-sided *t*-test. (**F**) Venn diagram and (**G**) heat map showing up-regulated genes in *met-2Δ hpl-2Δ* double mutant early embryos measured by RNAseq, and the subset of genes independently observed to be de-repressed in *hpl-2Δ* and/or *met-2Δ* single mutant early embryos. N = 3, FC > 2, FDR < 0.05. (**H**) qRT-PCR of a subset of *met-2Δ hpl-2Δ* double mutant up-regulated genes or the control gene *lmn-1*. Bars show means while dots indicate individual measurements. N = 3.

For a comprehensive view of endogenous gene expression, we performed RNAseq. We compared *met-2Δ* with *hpl-2Δ* mutant early embryos and found 1319 genes up-regulated in *met-2Δ* and 205 genes up-regulated in *hpl-2Δ* (Fig. 2C, FC = 2, FDR = 0.05). Remarkably, not many genes were shared between the two mutant backgrounds (4% of MET-2 and 27% of HPL-2 targets shared). Comparison of HPL-2 up-regulated genes with those of *set-25Δ* mutants, lacking all H3K9me3, showed a similar lack of overlap (Fig. S2A), as SET-25 represses the same genes as MET-2 (Fig. S2B). In mutants lacking H3K9me, gene activation is associated with increased histone acetylation (25). Western immunoblotting confirmed that H3K9ac and H3K27ac were upregulated in *met-2Δ* embryos, but we observed no change in acetylation of these H3 residues from WT in *hpl-2Δ* mutants (Figs. 2D-E). These data show that MET-2 and HPL-2 silence distinct cohorts of genes and have different effects on global acetylation levels.

We next tested whether *met-2Δ hpl-2Δ* double mutants function redundantly for transcriptional repression. We performed RNAseq on *met-2Δ hpl-2Δ* double mutant early embryos. Sterile double mutants are maternally rescued, so we were able to employ a fluidic sorting strategy to purify a population of fertile double mutants from which to isolate embryos (see Methods). We found 928 genes upregulated in *met-2Δ hpl-2Δ* embryos relative to WT (Fig. 2F-G, FC = 2, FDR 0.05). Of these 928 upregulated genes, 543 and 125 genes were previously seen upregulated in *met-2Δ* and *hpl-2Δ* single mutants, respectively. However, 352 (38.2%) genes were de-repressed only in double mutant embryos. We tested a subset of desilenced genes using qRT-PCR and confirmed synergistic upregulation in *met-2Δ hpl-2Δ* embryos (Fig. 2H), demonstrating that MET-2 and HPL-2 act redundantly to repress certain genes. We also observed certain repeat families were upregulated only in the double mutants (Fig. S2C). Interestingly, GO term analysis of genes upregulated only in the double mutants enriched most prominently for histone genes (Fig. S2D-E). Taken together, the data show that MET-2 and HPL-2 act redundantly to silence a subset of genes and repeats, suggesting MET-2 and HPL-2 function in independent pathways to silence some genes but synergize to silence others.

### HPL-2 subnuclear foci are reinforced, but largely independent of H3K9me

Since our transcriptional data suggested an H3K9me-independent function of HPL-2, we tested whether HPL-2 foci are in any way dependent on H9K9me. In *met-2Δ set-25Δ* double mutant embryos lacking all H3K9me, HPL-2 still formed foci (Fig. 3A). Therefore, H3K9me is not necessary for HPL-2 foci formation. This result was unexpected because existing models suggest H3K9me2/me3 binding would be a driving force in the local accumulation of HPL-2(9, 10). We therefore interrogated the properties of HPL-2 foci using automated image analysis in WT and H3K9me-deficient backgrounds. Despite HPL-2 retaining its foci-forming competence in the absence of H3K9me, foci were fewer in number (WT: mean 5.7+/- 0.6 and *met-2Δ set-25Δ*: mean 3.6 +/- 0.5 foci/nucleus; p-value: 0.017), had a reduced fluorescence intensity (WT: mean 4.6+/- 0.4 and *met-2Δ set-25Δ*: mean 2.9 +/- 0.3 a. u.; p-value: 0.0046), and were measurably smaller (WT: mean 29.0+/- 0.5 and *met-2Δ set-25Δ*: mean 24.7 +/- 0.6 voxel/foci; p-value: 0.0036) in *met-2Δ set-25Δ* double mutant embryos on average (Fig. 3B-D). These changes could not be attributed to less HPL-2 protein overall, as both fluorescence intensity and western immunoblotting showed no decrease of total HPL-2 protein signal in the absence of H3K9me (Fig. 3E-F). Therefore, HPL-2 form foci even in the absence of H3K9me, but H3K9me may promote foci stability and/or assembly. We also observed that HPL-2 foci accumulate at the nuclear periphery in WT embryos (Fig. 3G) (26). Consistent with previous studies of MET-2 foci and heterochromatin, HPL-2 foci shifted away from the periphery when H3K9me was lost in the *met-2Δ set-25Δ* mutants (26, 38–40). Therefore, HPL-2 foci are dependent on MET-2 and SET-25 for targeting to the nuclear periphery.

**Fig 3.**
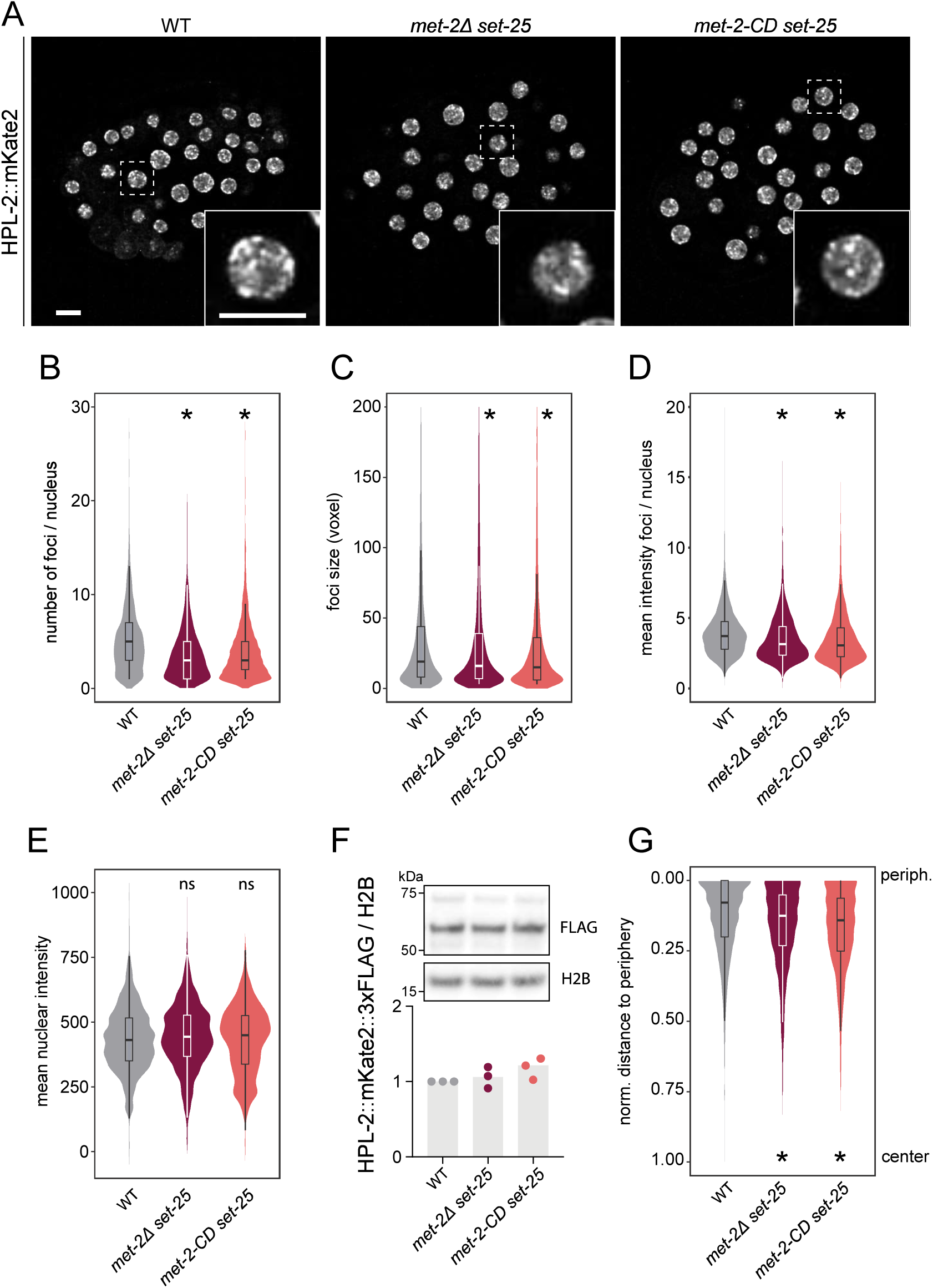
HPL-2 forms nuclear foci independent of H3K9 methylation. (**A**) Representative live images of HPL-2::mKate2::3xFLAG in WT, *met-2Δ set-25Δ*, or *met-2-CD set-25Δ* embryos. Scale bar, 5 μm. N = 3, n = 30. Quantification of HPL-2::mKate2 foci from (A) measuring (**B**) foci number per nucleus, (**C**) size of foci, (**D**) mean foci intensity per nucleus, and (**E**) mean nuclear intensity. (**F**) Representative western blot of HPL-2::mKate2::3xFLAG protein levels in WT, *met-2Δ set-25Δ*, or *met-2-CD set-25Δ* embryos. N = 4. (**G**) Quantification of HPL-2::mKate2 foci distance to the nuclear periphery. *P-values (*met-2Δ set-25Δ*, *met-2-CD set-25Δ* vs WT): (B) p = 0.017, p = 6.0 x 10^-3^, (C) p = 4.2 x 10^-3^, p < 3.6 x 10^-3^, (D) p = 3.1 x 10^-4^, p = 4.6 x 10^-3^, (E) not significant, (G) p = 2.1 x 10^-13^, p < 2.2 x 10^-16^, by one-way ANOVA, followed by Tukey post hoc test.

Given MET-2’s non-catalytic activity in heterochromatin function, we sought to distinguish whether H3K9me or MET-2 itself was important for HPL-2 foci or lamina association using catalytically deficient mutant *met-2-CD*. Animals expressing *met-2-CD; set-25Δ* did not rescue HPL-2 foci robustness (Fig. 3B-F), indicating that H3K9me rather than MET-2 reinforces HPL-2 foci. Likewise, *met-2-CD* could not rescue the association of HPL-2 with the nuclear lamina (Fig. 3G), implicating H3K9me in targeting HPL-2 foci to the lamina. In the reverse experiment, deletion of *hpl-2* (*hpl-*2*Δ*) did not decrease MET-2::mNeonGreen foci number, intensity, size, or enrichment at the periphery (Fig. S3A-E). Therefore, H3K9me reinforces HPL-2 protein foci and localization, while MET-2 traffics independently of HPL-2.

### HPL-2 defective in H3K9me-binding retains transcriptional repression and foci-forming competence

To test directly the role of methyl-lysine binding for HPL-2 foci formation, we disrupted HPL-2’s ability to bind H3K9me. HP1 binds H3K9me2/3 via a three-amino acid aromatic cage in the chromodomain (Fig. 4A). We mutated two phenylalanines of the HPL-2 aromatic cage to alanine, F19A and F43A (Fig. 3A, HPL-2-FFAA). Analogous mutations in other chromodomain proteins have been shown to disrupt H3K9me binding (39, 41). We confirmed that this HPL-2FFAA double point mutant showed strongly impaired interaction with H3K9me2/3 using an in vitro binding assay (Fig. S4A). Protein levels of mKate2-tagged strains were not decreased in the HPL-2-FFAA strain relative to WT, indicating that the mutation does not affect expression (Fig. S4B-C).

**Fig 4.**
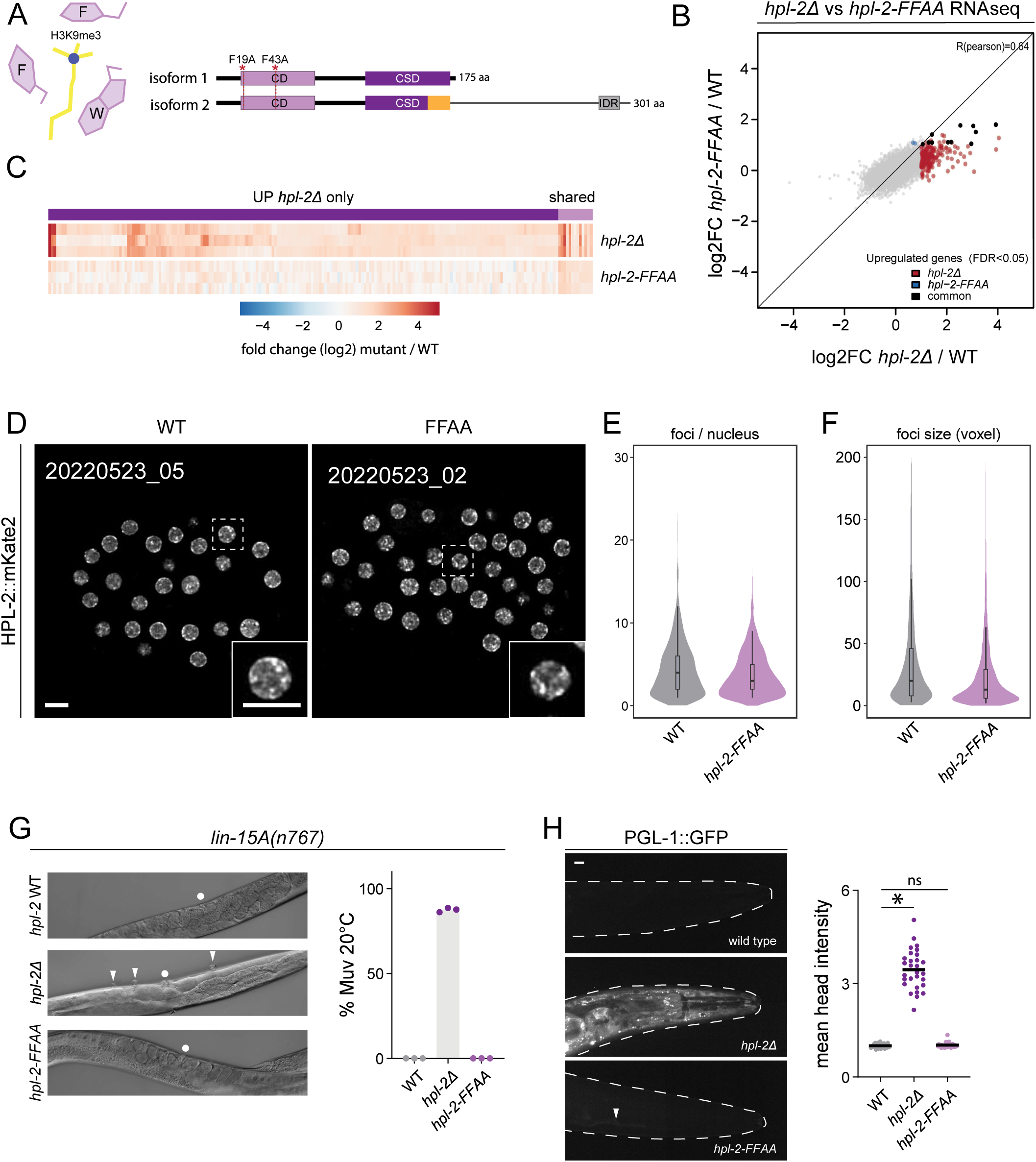
HPL-2 defective in binding H3K9me2/3 suffices for foci formation and cell identity. (**A**) Schematic of the HPL-2 aromatic cage (F19, W41, F43) (53, 92), and the *hpl-2-FFAA* double point mutation. (**B**) RNAseq scatterplot and gene heat map (**C**) comparing gene expression between *hpl-2Δ* and *hpl-2-FFAA* mutant embryos. N = 3, FC > 2, FDR < 0.05 (**D**) Representative live images of HPL-2::mKate2::3xFLAG in WT or *hpl-2-FFAA* embryos. Scale bar, 5 μm. N = 3, n = 30. Quantification of HPL-2::mKate2 foci (**E**) number and (**F**) size from data in (D). N = 3. *P-values (*hpl-2-FFAA* vs *hpl-2-WT*): (E) p = 2.8 x 10^-3^, (F) p = 2.8 x 10^−3^ by one-way ANOVA, followed by Tukey post hoc test. (**G**) Representative DIC live images and quantification of multivulva (Muv) defects in worms expressing *hpl-2-WT*, *hpl-2Δ* or *hpl-2-FFAA* in the sensitized *lin-15A(n767)* background. Normal vulvae = circle, multivulva defects = arrowheads. N = 3. n: WT = 1437, *hpl-2Δ* = 1975, *hpl-2-FFAA* = 1483. (**H**) Representative live image maximum projections and signal quantification of endogenously-tagged PGL-1::GFP in the heads of L4 animals. Body is outlined in dashed line; mouth is to the right. Arrowhead = faint ectopic expression. Scale bar = 10 μm, N = 3, n = 30. *p < 10^-5^, ns = not significant by one-way ANOVA, followed by Tukey post hoc test.

If HPL-2 indeed has H3K9me-independent functions in ensuring transcriptional silencing, then the *hpl-2-FFAA* mutant may maintain gene repression of at least a subset of genes mis-expressed in the *hpl-2* deletion mutant. We therefore performed transcriptome analysis comparing WT, *hpl-*2*Δ*, and *hpl-2-FFAA* early embryos. We found that genes desilenced in *hpl-*2*Δ* embryos remained largely repressed in *hpl-2-FFAA* animals (Fig. 4B-C, S4D; FC = 2, FDR = 0.05), indicating that HPL-2 can silence genes in an H3K9me-independent manner. Nevertheless, gene expression in *hpl-2-FFAA* and *hpl-2Δ* correlated (R = 0.64), suggesting that the repression was not complete. We conclude that HPL-2 possesses a silencing function that does not require binding H3K9me.

Despite its compromised ability to bind H3K9me, HPL-2-FFAA mKate2-tagged protein formed foci (Fig. 4D). Consistent with the properties of HPL-2 foci in the *met-2Δ set-25Δ* double mutant lacking all H3K9me, the number and average size of foci were decreased in HPL-2-FFAA relative to HPL-2-WT (Fig 4D-F). These results demonstrate a direct role for H3K9me-binding in reinforcing the assembly or stability of HPL-2 foci. In contrast, foci intensity and peripheral positioning showed only modest effects in HPL-2-FFAA mutants vs. WT embryos (Fig. 4D, S4E-F), suggesting that H3K9me may exert some effects on HPL-2 protein indirectly. Further, HPL-2-FFAA continued to colocalize with MET-2-WT and MET-2-CD (Fig. S4G), suggesting that HPL-2-FFAA foci are correctly targeted despite its H3K9me-binding deficiency.

Having uncoupled HPL-2 protein foci from H3K9me binding, we tested whether HPL-2-FFAA could perform known functions of HPL-2. We first compared the ability of *hpl-2Δ* and *hpl-2-ffaa* mutants to promote proper tissue differentiation. Both HPL-2 and MET-2 function in the development of the adult *C elegans* vulva (21). *hpl-2* and *met-2* are classified into the genetically defined category of SynMuvB genes, which are typically thought of as a single pathway. Single mutant animals develop a normal vulva. In the presence of an additional mutation in a SynMuvA gene, such as *lin-15A*, doubly mutant animals fail to restrict vulval development in the maturing epidermis, resulting in ectopic pseudovulvae, also known as a <u>Mu</u>lti<u>v</u>ulva phenotype (Muv). We crossed *hpl-*2*Δ* or *hpl-2-FFAA* animals into a *lin-15A* mutant background. As expected, *hpl-*2*Δ;lin-15A* animals showed a high incidence of multivulva (Fig. 4G). In contrast, *hpl-*2-*FFAA; lin-15A* animals showed normal vulva development. Similar to *hpl-2-FFAA* animals, catalytically deficient *met-2-CD; lin-15A* animals also developed normally, while *met-2Δ; lin-15A* animals displayed a highly penetrant multivulva phenotype (Fig. S4H).

These data show that both HPL-2 and MET-2 can preserve normal vulval development even when their H3K9me-related functions are compromised.

Animals with mutations in the SynMuvB heterochromatin genes are also known to ectopically express germline genes in somatic cells (42–45). We asked whether this soma-to-germline phenotype was H3K9me-dependent using an endogenous GFP-tagged P-granule protein PGL-1::GFP (32, 46). We imaged the heads of L4 larvae to avoid confounding autofluorescence in the gut and the normal PGL-1 signal in the germline. While deletions of either *met-2* or *hpl-2* lead to a several-fold increase in ectopic GFP expression in the heads, animals expressing *met-2-CD* or *hpl-2-FFAA* retained nearly complete repression of PGL-1 expression (Figs. 4H, S4I). Taken together, these results show that both HPL-2 and MET-2 have H3K9me-independent functions in foci formation, transcriptional repression and vulva cell fate.

### Physical interaction with multivalent binding partners are essential for MET-2 and HPL-2 H3K9me independent functions

Because both HPL-2 and MET-2 exhibit H3K9me-independent functions, we examined how their interaction partners contribute to foci formation and function. MET-2 foci are dependent on physical interaction with the intrinsically disordered domain (IDR) protein LIN-65 (26, 27). In the absence of *lin-65*, MET-2 does not concentrate in the nucleus and does not form foci. Similarly, an overexpressed HPL-2 was reported to form foci dependent on the C2-H2 zinc finger-containing protein LIN-13 (29). LIN-65’s IDRs and LIN-13’s predicted 24 zinc fingers create the potential for higher order interactions. Such multivalent interactions have been observed to contribute to biomolecular condensates (47, 48), and indeed MET-2 foci possess some properties associated with condensates (26). We interrogated whether MET-2 and/or HPL-2 foci are dependent on either or both of their multivalent co-factors LIN-65 and/or LIN-13, respectively. Knockdown of *lin-65* dispersed MET-2 from the nucleus as expected; however, HPL-2 foci remained (Fig. 5A). Knockdown of *lin-13* depleted HPL-2 foci as previously reported but was dispensable for MET-2 foci formation (Fig. 5A). We conclude HPL-2 and MET-2 colocalize in foci independent of each other or the other’s required cofactor, LIN-13 and LIN-65, respectively.

**Fig. 5.**
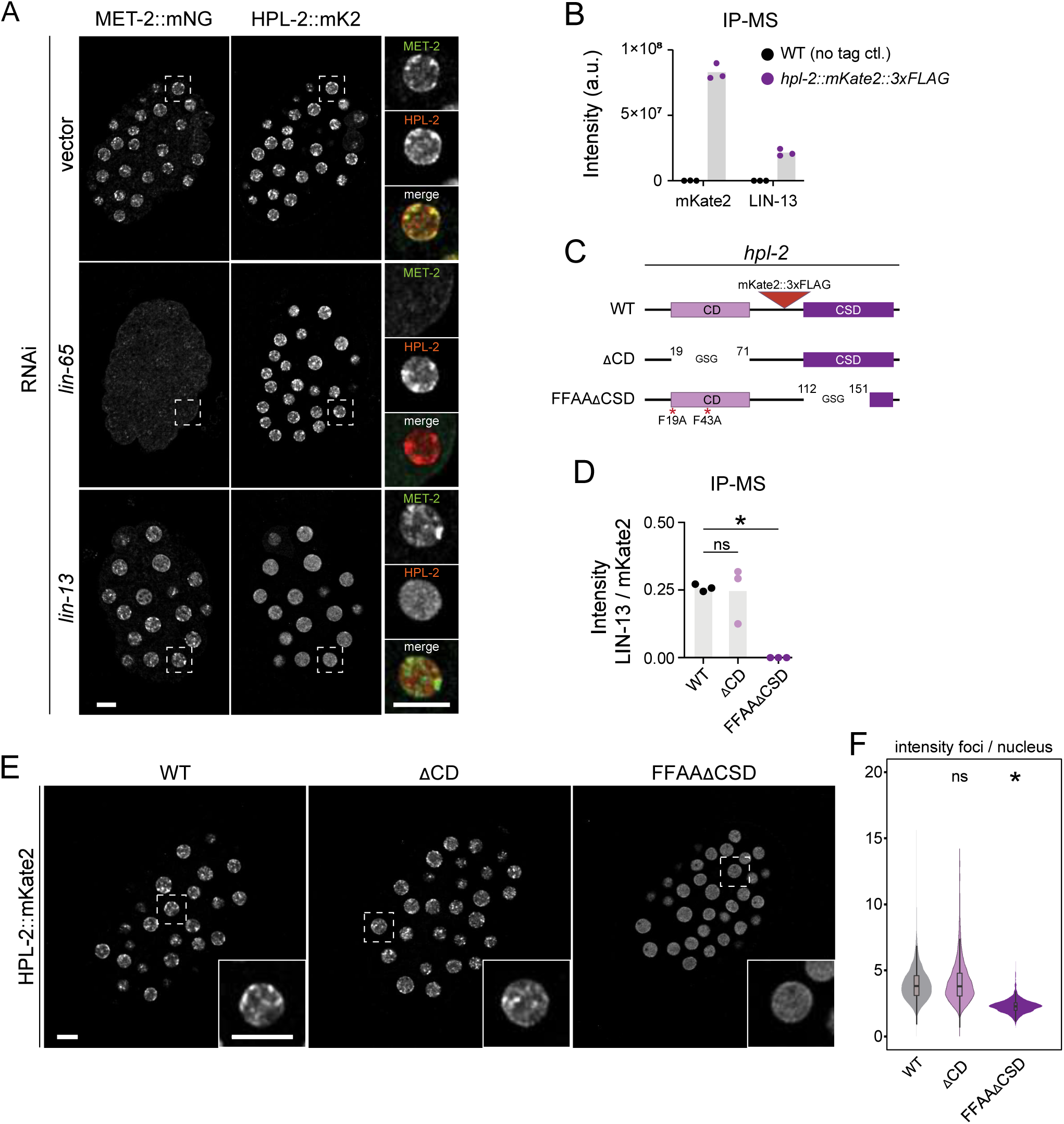
HPL-2 physical interactions via its chromoshadow domain promote foci formation. (**A**) Representative images of embryos co-expressing MET-2::mNeonGreen with HPL-2::mKate2 treated with the indicated RNAi or empty vector control. Scale bar, 5 μm. N = 3, n = 30. (**B**) anti-FLAG immunoprecipitation coupled to mass spectrometry enrichment of mKate2 and LIN-13 in untagged and HPL-2::mKate2::3xFLAG embryos. N = 3. (**C**) Schematic of chromodomain (ΔCD) or chromoshadow domain (ΔCSD) deletions created in HPL-2::mKate2 expressing animals (WT or FFAA, respectively). Positions of first and last amino acids of the deletions, replaced with Gly-Ser-Gly, are indicated. For clarity, only the short isoform is shown. (**D**) LIN-13 spectral intensity normalized to that of mKate2 in WT, *hpl-2-ΔCD*, and *hpl-2-FFAAΔCSD*. N = 3. *p = 4 x 10^-6^, ns = not significant by two-sided T test. (**E**) Representative live images of HPL-2::mKate2 in WT, HPL-2-ΔCD, and HPL-2-FFAAΔCSD embryos. Scale bar, 5 μm. N = 3, n = 30. (**F**) Quantification of foci intensity relative to nuclear signal from (E). HPL-2ΔCD vs WT not significant, HPL-2-FFAAΔCSD vs WT p < 2.2 x 10^-16^ using one-way ANOVA, followed by Tukey post hoc test.

We asked how LIN-13 and LIN-65 contribute to HPL-2 and MET-2 function in cell fate. The catalytic and non-catalytic functions of MET-2 are dependent on LIN-65 (25). Conversely, we also observed that LIN-65::GFP was lost in *met-2Δ* nuclei (Fig. S5A). Adding an NLS-tag to the C-terminus of LIN-65::GFP did not rescue LIN-65 in a *met-2*-deficient background, suggesting that LIN-65 is degraded rather than delocalized (data not shown). In contrast, *met-2-CD* rescued LIN-65::GFP nuclear foci (Fig. S5A, S5B). These results show that MET-2 and LIN-65 are mutually dependent for stable accumulation into nuclear foci, but independent of H3K9me. Like MET-2, deletion of *lin-65* shows a high penetrance multivulva phenotype in the *lin-15A* mutant context (Fig. S5C and (49)). While loss of *lin-65* disrupts H3K9me, MET-2 nuclear accumulation and foci formation, loss of MET-2’s catalytic cofactor ARLE-14 disrupts H3K9me deposition and MET-2 chromatin association without disrupting MET-2 nuclear foci (25, 27). We found that vulval development was normal in *arle-14Δ;lin-15A* double mutant animals (Fig. S5C). These results are consistent with an H3K9me-independent mechanism for vulval development. Next, we tested whether nuclear localization in the absence of *lin-65* was sufficient for MET-2 to promote cell fate by expressing MET-2 with a nuclear localization sequence (NLS; (27)). It was previously shown that this NLS could maintain MET-2 nuclear enrichment in the absence of *lin-65*; however, neither nuclear foci nor H3K9me levels were rescued (25). Just like H3K9me levels, *NLS::met-2* was not sufficient for normal vulval development in the absence of *lin-65* (Fig. S5D). Fertility assays also showed no rescue of progeny numbers in the *lin-65Δ;NLS::met-2* double mutant animals (Fig. S5E). Taken together, these results suggest that LIN-65 is stabilized by interaction with MET-2 and has an essential role in cell fate, correlating with foci formation, that cannot be rescued by either MET-2 catalytic activity or MET-2 accumulation in the nucleus.

Next, we examined the interaction between HPL-2 and LIN-13. A previous report demonstrated that HPL-2 and LIN-13 directly bind each other in yeast two hybrid studies (29). We performed immunoprecipitation of HPL-2::mKate2::3xFLAG followed by mass spectrometry (IP-MS) to characterize HPL-2 interactions in embryo lysates. LIN-13 peptide intensities were indeed enriched in the HPL-2::mKate2 strain relative to the N2 untagged control (Fig. 5B). Consistent with a model of independent recruitment, HPL-2 pulldowns did not enrich for MET-2, nor did previous MET-2 pulldowns enrich for HPL-2 (data not shown and (25–27).

We characterized in greater detail how the physical interaction with LIN-13 contributes to HPL-2 function. The data above demonstrating that the HPL-2-FFAA mutant suffices for vulval development and transcriptional repression argues against an essential role for the chromodomain. In contrast, two-hybrid interaction studies suggest that the HPL-2 chromoshadow domain (CSD) suffices for binding LIN-13, which is also implicated in proper vulva specification (29, 50–52). To test which domains were necessary for HPL-2 function, we used CRISPR to delete HPL-2’s chromodomain (aa 19-71, ΔCD) or the portion of the CSD shared between both short and long isoforms that contains the conserved residues necessary for binding the PXVXL motif (53) in the FFAA background (aa 112-151, ΔCSD) (Fig. 5C). To characterized these domain mutants, we examined the interaction with LIN-13 via immunoprecipitation of the HPL-2 mutants followed by mass spectrometry (IP-MS). While LIN-13 remained enriched in ΔCD mutants to a similar extent as WT, LIN-13 enrichment was lost in FFAAΔCSD embryos (Fig. 5D). Therefore, the HPL-2 chromoshadow domain is necessary for interaction with LIN-13 in vivo. Consistent with these data, the HPL-2 signal enriched in foci was retained in embryos lacking the chromodomain but lost in ΔCSD embryos, confirming that LIN-13-dependent physical interaction mediated by the chromoshadow domain are necessary for foci formation (Fig. 5E, 5F).

We tested how HPL-2 domains interacted with *lin-15A* in specification of the vulva. *hpl-2ΔCD;lin-15A* animals showed a low penetrance multivulva phenotype (8.2%, Fig. 6A). It follows we cannot exclude a structural role for the chromodomain in this process. In contrast, animals lacking the chromoshadow domain showed highly penetrant defects in vulva development (90.9%, Fig. 6A). Finally, we used qRT-PCR to interrogate genes identified by RNAseq to be repressed by HPL-2. Deletion of the chromodomain showed only modest effects on transcription, while embryos lacking the chromoshadow domain exhibited greater desilencing of genes, similar in magnitude to embryos in which *hpl-2* is deleted (Fig. 6B). Therefore, we conclude the HPL-2 chromoshadow domain mediates interactions with its multivalent binding partner LIN-13 to drive foci formation, repress transcription, and promote proper tissue differentiation.

**Figure 6.**
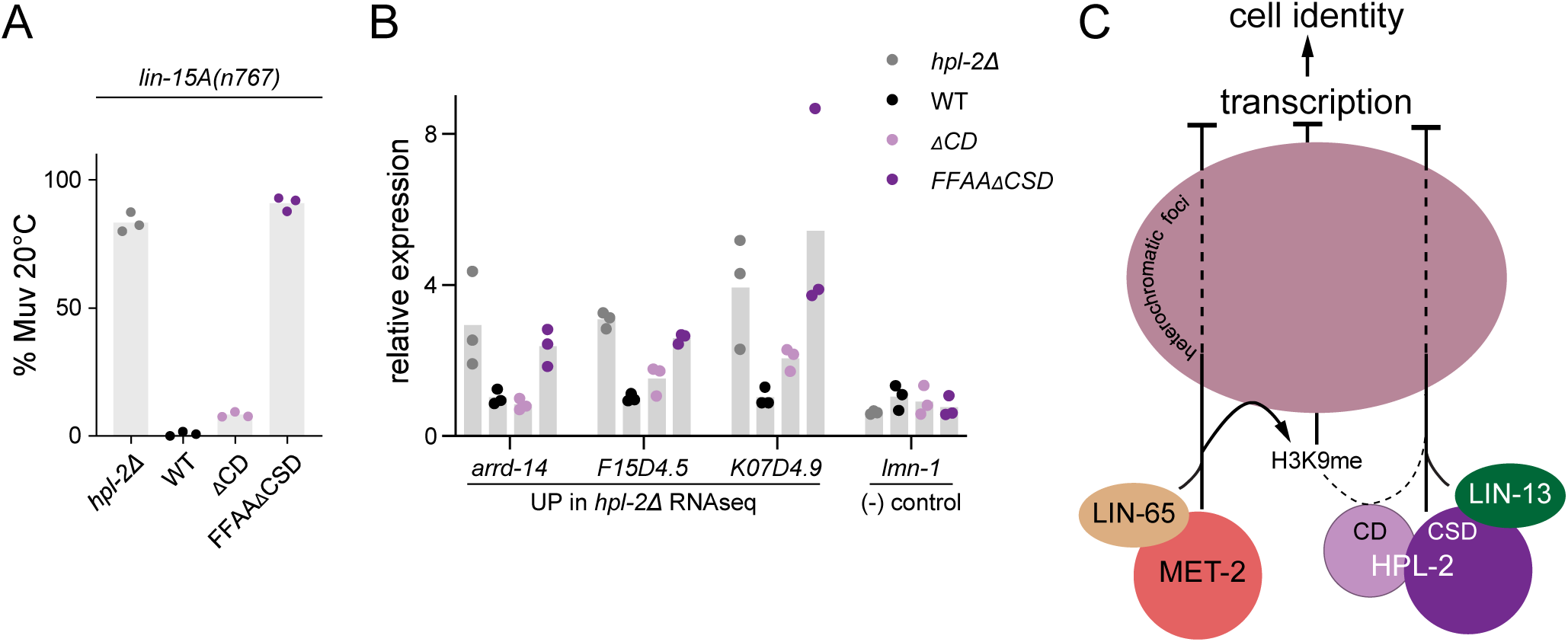
HPL-2 chromoshadow domain is essential for cell fate acquisition. (**A**) Quantification of multivulva (Muv) defects in worms expressing, *hpl-2Δ* or mKate2-tagged strains shown in (A) in the sensitized *lin-15A(n767)* background. N = 3. n: *hpl-2Δ* = 859, WT = 899, *hpl-2-ΔCD* = 887, *hpl-2-FFAAΔCSD* = 941. (**B**) qRT-PCR of a subset of *hpl-2Δ* up-regulated genes or the control gene *lmn-1* in *hpl-2Δ* or mKate2-tagged strains shown in (A). Bars show means while dots indicate individual measurements. N = 3. (**C**) Model: Interactions with multivalent cofactors are necessary for H3K9me-indpendent functions of HPL-2 and MET-2 during development. CSD-dependent protein-protein interactions with LIN-13 recruits HPL-2 into foci and suffices to restrict cell fate and repress transcription while MET-2 catalytic and non-catalytic activities require the intrinsically disordered domain protein LIN-65.

## Discussion

Recent studies have shown that multiple histone-modifying enzymes retain function in the absence of their enzymatic activity, raising the question of how these proteins interact with the protein readers of histone modifications in the absence of enzymatic activity. We find that the reader HPL-2/HP1 and writer MET-2/SETDB1 function in parallel pathways to silence distinct groups of genes (Fig. 6C). They also act redundantly and/or additively to silence a second cohort of sequences. These activities depend on physical interactions with distinct multivalent binding partners, the zinc finger LIN-13 for HPL-2 and the intrinsically disordered LIN-65/ATF7IP for MET-2. Interactions with these cofactors are critical for gene repression, organogenesis and localization of HPL-2 and MET-2 within heterochromatic foci. H3K9me is not necessary for either factor to concentrate into foci, but does contribute to HPL-2 foci robustness and distribution. These protein-protein interactions may explain the minor role of H3K9me binding in HPL-2-mediated gene repression.

The histone code hypothesis argues that a combinatorial crosstalk between post-translational modifications of histones regulates chromatin function, including the selective expression of genes (54). In particular, the role of H3K9 methylation in silencing of genes and repetitive elements is well-established (19, 55–58). Nevertheless, increasing evidence is uncoupling PTM writer proteins from their histone marks in their regulation of transcription and development. In Drosophila, the histone acetyltransferases (HATs) CBP and Gcn5 are required for zygotic genome activation, yet can perform their essential role in development independent of its acetylation activity (59). In mammalian cells, p300 plays an important role in transcription coactivation, but chromatin interactions have been reported to be opposed by its HAT activity (60). Use of chemical inhibitors of HAT activity has strengthened the evidence for acetylation-independent functions of CBP (61). Among methyltransferases, mammalian MLL3 and MLL4 have been shown to promote enhancer activation independent of their H3K4 HMT activity (62). G9a can function in transcriptional repression independent of its H3K9 HMT activity by recruiting DNA methyltransferases (63, 64). We found that the catalytic-deficient MET-2-CD rescues or mitigates germline and somatic phenotypes, prevents hyperacetylation, and is sufficient to silence a subset of genes repressed by MET-2 (Fig. S4H-I, and (25)). Intriguingly, a subsequent study showed SETDB1 bearing the homologous C1243A catalytic deficient mutation as MET-2-CD rescues defects in stem cell pluripotency caused by simultaneous triple-deletion of HP1α/β/γ, arguing that a non-catalytic activity of MET-2/SETDB1 may be conserved (65). MET-2-CD stabilizes LIN-65/ATF7IP, and both proteins are required for cell fate restriction (Fig S5). These findings suggest that protein-protein interactions between chromatin modifiers and their cofactors are required for an additional layer of regulation beyond that mediated by PTMs.

PTM reader proteins can also demonstrate dynamic, higher-order protein behaviors that are enhanced by, but not dependent on, chromatin marks. It has been shown, for example, that histone acetylation reader BRD4 forms condensates via its IDR, but that biophysical properties of these foci are altered in the presence of acetylation (18, 66, 67). We show that HPL-2 need not bind to H3K9me to repress transcription or promote appropriate vulva cell lineage; rather, H3K9me contributes to robust HPL-2 gene repression and protein dynamics in vivo. Consistently, in flies H3K9me enhances, but is not required for, HP1a accumulation at promoters and pericentromeric regions (68, 69). In contrast, H3K9me is required for anchoring of HPL-2 foci to the periphery (Fig. 3G). HP1 proteins consist of a H3K9me-reader chromodomain, a flexible hinge region, and a chromoshadow domain that conveys homo- and heterodimerization with binding partners. In mammals, the chromoshadow domain leads to the recruitment of Suvar3-9 and Suv420H2, leading to more repressive H3K9me and H4K20me3, respectively (70, 71). Several CSD interactions with chromatin-associated factors have been reported, including CAF-1, a nucleosome chaperone, and KAP-1, which forms a complex with SETDB1 and also associates with KRAB zinc finger proteins (72–76) . We demonstrated the CSD is required for interaction between HPL-2 and LIN-13 in vivo, the concentration of HPL-2 into foci, and the specification of the mature vulva. In contrast, the chromodomain played only a minor role in cell fate, transcription, and protein foci dynamics.

Multiple mechanisms may underlie how multivalent interactions concentrate proteins into functional compartments. For example, it has been proposed that p300 acts as a scaffold, activating transcription by simultaneously recruiting multiple transcription factors (60). Intrinsically disordered domains of proteins (IDRs) are strongly implicated in higher-order protein condensation (77). IDRs are unstructured but have unique biochemical attributes including electrostatic charge and primary sequence that have been shown to control protein targeting and function (78, 79). Post translation modification of IDRs have also been shown to regulate their function. Phosphorylation of HP1α unstructured regions was demonstrated to enhance intermolecular self-interactions, lowering the barrier to phase separation in vitro (17, 80). Our data and others demonstrate that HP1 homolog HPL-2 forms foci in vivo not via H3K9me or self-interactions but by binding LIN-13, whose 24 zinc fingers and LIN-35/Rb binding motif may serve as a scaffold promoting HPL-2 multivalent protein-protein and protein-chromatin interactions (29). Analogously, MET-2/SETDB1 possesses an unstructured N-terminus that is not sufficient to establish and/or maintain foci; rather, MET-2, likely via its N-terminus, interacts with the disordered protein LIN-65/ATF7IP to be recruited into higher-order structures, methylate H3K9, and repress transcription (25–28, 81–83). Importantly, the interplay between HPL-2 and MET-2 with their respective putative multivalent cofactors do not only drive foci formation, but together they provide an additional layer of regulation essential for cell lineage restriction and the silencing of germline genes in the soma (42–45). LIN-13, MET-2, and HPL-2 have been suggested to work together with other heterochromatin-associated proteins to prevent DNA damage and germline apoptosis (19). We find that MET-2 and HPL-2 colocalize in foci in a LIN-13 dependent manner. Dissecting to what extent multivalent protein-protein and protein-chromatin interactions contribute to PTM-independent regulation of transcription and tissue identity remains an important challenge worthy of subsequent investigation.

## Materials and Methods

### Strains and maintenance

Strains used for these studies are documented in Supplementary Table 1. “Some strains were provided by the CGC, which is funded by NIH Office of Research Infrastructure Programs (P40 OD010440).” Animals were cultured at 20°C unless otherwise indicated. Embryos were isolated either by dissection or by standard bleaching protocols. RNA interference (RNAi) knockdown of genes was achieved as previously described (84, 85). Synchronized L1 animals were fed HT115 E coli expressing double stranded RNA targeting the indicated gene on nematode growth media plus 1 mM IPTG and 100 μg/mL carbenicillin. RNAi targeting *lin-13* was a gift from Francesca Palladino (CNRS Lyon).

### Transgenesis

Targeted substitutions and deletions were created in endogenous gene loci using CRISPR-Cas9 exactly as previously described (86). AltR Cas9 and tracrRNA (IDT) were used. AltR crRNAs were synthesized by IDT. Edit-specific crRNA sequences and Homology Directed Repair template sequences (IDT) are provided in Supplementary Table 2.

### Live microscopy

Imaging was performed at 20°C. 2D DIC images were captured using a Zeiss Axio Zoom V16 and Zeiss Axiocam 503 mono camera with ZEN 2.6 blue edition software. Fluorescent 3D imaging were obtained using an Olympus IX83 microscope outfitted with a Yokogawa CSU-W1 confocal 50 μm spinning disk, and Hamamatsu ORCA-Fusion sCMOS camera. Images were deconvolved with the Huygens Remote Manager. Automated image segmentation analysis of nuclei and foci was performed with Knime (87).

### Developmental phenotypes

#### Fertility assays

L4 animals grown at the indicated temperature were placed one (25°C) or three (26°C) animals per plate. Egglayers were transferred to fresh plates every 24 hr for the duration of egglaying, after which total progeny were counted.

#### SynMuv assays

Gravid adults were allowed to lay eggs for several hours, then removed. Laid eggs were allowed to mature to day 1 adults for 72 hr at 20°C, at which time animals were visually scored for the presence or absence of abnormal vulva development.

### Immunoprecipitation-mass spectrometry (IP-MS)

Immunoprecipitations were performed as previously reported (26). All steps were performed on ice or 4°C. Briefly, mixed embryos isolated by bleaching from 100,000 day 1 adult animals were homogenized in TAP buffer (150 mM NaCl, 20 mM Tris-HCl, pH 7.5, 0.5% NP-40, 1 mM EDTA, 10% glycerol, 2× cOmplete-EDTA-free protease inhibitors (Roche), 1× Deacetylase Inhibitor (Active Motif), and 1 mM DTT) using 0.5 mm Glass Globules (Roth) and a Fast Prep-24 homogenizer (MP Biomedicals). Lysates were sonicated in 14 cycles of 15 s on, 30 s off using a Bioruptor Plus (Diagenode). Lysates were cleared via centrifugation at 21,000g for 10 min at 4°C. Anti-FLAG-M2 magnetic beads (Sigma) pre-washed in TAP buffer were added to 5 mg protein lysate and rotated overnight at 4°C. Beads were washed in TAP buffer 4 x 15 min, then 3 times with MS wash buffer (20 mM Tris-HCl, pH 7.5, 150 mM NaCl) to remove detergents.

Resin was resuspended in elution buffer (5% SDS, 10mM TCEP, 0.1 M TEAB), incubated for 10 min at 95°C and transferred into a new tube. Proteins were alkylated in 20 mM iodoacetamide for 30 min at 25°C and digested using S-Trap™ micro spin columns (Protifi) according to the manufacturer’s instructions. Briefly, 12 % phosphoric acid was added to each sample (final concentration of phosphoric acid 1.2%) followed by the addition of S-trap buffer (90% methanol, 100 mM TEAB pH 7.1) at a ratio of 6:1. Samples were mixed by vortexing and loaded onto S-trap columns by centrifugation at 4000 g for 1 min followed by three washes with S-trap buffer. Digestion buffer (50 mM TEAB pH 8.0) containing sequencing-grade modified trypsin (1/25, w/w; Promega) was added to the S-trap column and incubate for 1h at 47 °C. Peptides were eluted by the consecutive addition and collection by centrifugation at 4000 g for 1 min of 40 ul digestion buffer, 40 μL of 0.2% formic acid and finally 35 μL 50% acetonitrile, 0.2% formic acid. Samples were dried using a Speedvac and stored at -20 °C until further use.

Dried peptides were resuspended in 0.1% aqueous formic acid and subjected to LC–MS/MS analysis using a Orbitrap Fusion Lumos Mass Spectrometer fitted with an EASY-nLC 1200 (Thermo Fisher Scientific) and a custom-made column heater set to 60°C. Peptides were resolved using a RP-HPLC column (75μm × 36cm) packed in-house with C18 resin (ReproSil-Pur C18–AQ, 1.9 μm resin; Dr. Maisch GmbH) at a flow rate of 0.2 μL/min. The following gradient was used for peptide separation: from 5% B to 12% B over 5 min to 35% B over 40 min to 50% B over 15 min to 95% B over 2 min followed by 18 min at 95% B. Buffer A was 0.1% formic acid in water and buffer B was 80% acetonitrile, 0.1% formic acid in water.

The mass spectrometer was operated in DDA mode with a cycle time of 3 seconds between master scans. Each master scan was acquired in the Orbitrap at a resolution of 120,000 FWHM (at 200 m/z) and a scan range from 375 to 1600 m/z followed by MS2 scans of the most intense precursors in the linear ion trap at “Rapid” scan rate with isolation width of the quadrupole set to 1.4 m/z. Maximum ion injection time was set to 50ms (MS1) and 35 ms (MS2) with an AGC target set to 1e6 and 1e4, respectively. Only peptides with charge state 2 – 5 were included in the analysis. Monoisotopic precursor selection (MIPS) was set to Peptide, and the Intensity Threshold was set to 5e3. Peptides were fragmented by HCD (Higher-energy collisional dissociation) with collision energy set to 35%, and one microscan was acquired for each spectrum. The dynamic exclusion duration was set to 30s.

The acquired raw-files were searched using MSFragger (v. 4.0) implemented in FragPipe (v. 21.1) against a C. elegans database (consisting of 26584 protein sequences downloaded from Uniprot on 20220222) and 392 commonly observed contaminants spiked with the sequence of mKate2 using the “LFQ-MBR” workflow. Quantitative data was exported from FragPipe and analyzed using the MSstats R package v.4.13.0. (https://doi.org/10.1093/bioinformatics/btu305). Data was normalized, imputed using “AFT model-based imputation” and q-values for pairwise comparisons were calculated as implemented in MSstats.

### in vitro binding assay

Embryo lysed in IVB buffer (30mM Tris-HCl pH 7.5, 600 mM NaCl, 1 mM EDTA pH 8.0, 5% glycerol, 1% Triton X-100, 2X cOmplete-EDTA-free protease inhibitors (Roche), and 1 mM DTT) (88) were processed exactly as in immunoprecitation above. Dynabeads™ MyOne™ Streptavidin C1 (Invitrogen) were rotated with 4X excess biotinylated H3 tail peptides (Anaspec) in TAP buffer for 4 hr at 4°C. Beads were then washed 3X in IVB buffer and stored on ice prior to use. Equivalent amounts of protein were distributed to tubes containing peptide-bead complexes and rotated overnight at 4°C. Beads were washed 4X with IVB buffer, then resuspended in TAP buffer and stored at -20°C until use.

### Western

10 μg of total protein from embryo lysates was separated on Criterion XT Bis-Tris gels (Bio-Rad) using either MES or MOPS buffer (Invitrogen). Proteins were transferred using a Trans-blot semi-dry system and Trans-blot Turbo Midi 0.2 µm PVDF pre-cut membranes (Bio-Rad). Membranes were blocked in either PBS + 0.5% Tween-20 (PBST) containing 5% milk (Sigma) or Protein-free blocking buffer (Pierce) depending on the antibody of interest. Primary antibodies include 1:10,000 mouse anti-H3K9ac and 1:10,000 mouse anti-H3K27ac (gifts of H. Kimura), 1:10,000 rabbit anti-H2B (Abcam ab1790), and 1:25,000-1:50,000 mouse anti-FLAG M2 conjugated to HRP (Sigma). Blots were rotated overnight at 4°C, washed three times in PBST, blocked again and exposed to HRP-conjugated secondary antibodies, when applicable, for 1 h at RT. Secondary antibodies include 1:100,000 goat anti-mouse IgG HRP (Jackson ImmunoResearch 115-035-146) or 1:100,000 goat rabbit IgG HRP (Jackson ImmunoResearch 111-035-144). We used Immobilon HRP substrate (Millipore) and detected signal using a Fusion FX camera operated with Fusion FX7 Edge Software (Vilber).

### RNA isolation

RNA was extracted from early embryos from synchronized cultures grown at 20°C as reported previously using Trizol (Invitrogen) (24). For in *met-2Δ hpl-2Δ* double mutant embryos, heterozygotes with a GFP + balancer were expanded. Non-fluorescent L1 progeny—maternally rescued, homozygous double mutant animals—were selected using a COPAS Biosort (Union Biometrica), from which early embryos were harvested. Embryos were freeze cracked five times, then RNA was extracted with choloroform followed by isopropanol precipitation. Further purification was performed with the RNA Clean and Concentrator kit (Zymo).

### qPCR

cDNA was synthesized using Maxima H Minus Reverse Transcriptase (ThermoFisher) according to manufacturer’s instructions. 1 ug RNA was primed using Oligo(dT)20 primer (Invitrogen). Reactions were performed using the PowerUp SYBR Green Master Mix (Applied Biosystems). Ct values were determined with Mastercycler EP realplex software. Housekeeping gene pmp-3 was used for normalization (89) and relative expression was calculated with the ΔΔCt method (90).

### RNA sequencing

Libraries were produced using the Stranded Total RNA Prep Ligation with Ribo-Zero Plus Kit (Illumina). rRNA was depleted using the Ribo-Zero Plus kit supplemented with custom made C. elegans specific rDNA oligos (IDT). Libraries were profiled in a DNA 1000 Chip on a 2100 Bioanalyser (Agilent technologies) and quantified using the Qubit 1x dsDNA HS Assay Kit, in a Qubit 4.0 Fluorometer (Life technologies). Equimolar amounts of indexed libraries were pooled and sequenced on a NextSeq 2000 (Illumina). Reads were analysed as described previously (25). Adapter were trimmed using Trimmomatic v0.39. Reads were aligned to the C. elegans genome (ce10) with the R package QuasR v1.44.0, (www.bioconductor.org/packages/2.12/bioc/html/QuasR.html). The command “proj <-qAlign(“samples.txt”,“BSgenome. Celegans.UCSC.ce10”, splicedAlignment=TRUE)” instructs hisat296 to align using default parameters, considering unique reads for genes and genome wide distribution. Count Tables of reads mapping within annotated exons in WormBase (WS220) were constructed using the qCount function of the QuasR package to quantify the number of reads in each window (qCount(proj, GRange_object, orientation=“same”)) and normalized by division by the total number of reads in each library and multiplied by the average library size. Transformation into log2 space was performed after the addition of a pseudocount of 12 to minimize large changes in abundance fold change (FC) caused by low count numbers. The EdgeR package v4.2 was applied to select genes with differential transcript abundances between indicated genotypes (contrasts) based on false discovery rates (FDR) for genes (FDR <0.05 and log2 fold change >1 for up regulated genes and <-1 for down regulated genes). Replica correlations are shown in Fig SXXX of GO-Terms were extracted from wormbase.org and the GO-Term and KEGG pathway enrichment analysis was performed using the gprofiler2 package (v 0.2.3) as an interface to g: Profiler (91).

## Acknowledgements

We would like to thank the Imaging Core facility (IMCF Biozentrum), the Proteomics core facility members (Biozentrum, University of Basel) and the Genomics Core Facility (IMB, Mainz) for helpful discussions and technical assistance; Francesca Palladino for clones, and Iskra Katic at Fredrich Miescher Institute (FMI) for providing sound advice and technical support for fluidic sorting of animals. Jan Padeken & Lukas Stelzl are supported by the SFB 1551 Project No. 464588647 of the DFG (Deutsche Forschungsgemeinschaft). SMG thanks the ERC AdvGrant Epiherigans. SEM is supported by Univ of Basel and SNF 10001070. CD and SEM were supported by the Novartis Stiftüng für medizinischbiologishe Forschung. Some strains were provided by the CGC, which is funded by NIH Office of Research Infrastructure Programs (P40 OD010440).

## Authors’ Contributions

CED: Conceived the project, performed experiments, data curation, formal analysis, methodology, Investigation: data in figures, Writing: original draft, review, editing.

JEY: Investigation: performed experiments

SG: Funding acquisition

JP: Investigation: data in figures; Writing: review, editing; funding acquisition

SEM: conceptualization; Writing: review, editing, supervision; funding acquisition, project administration

## Availability of data and materials

**Fig. S1.**
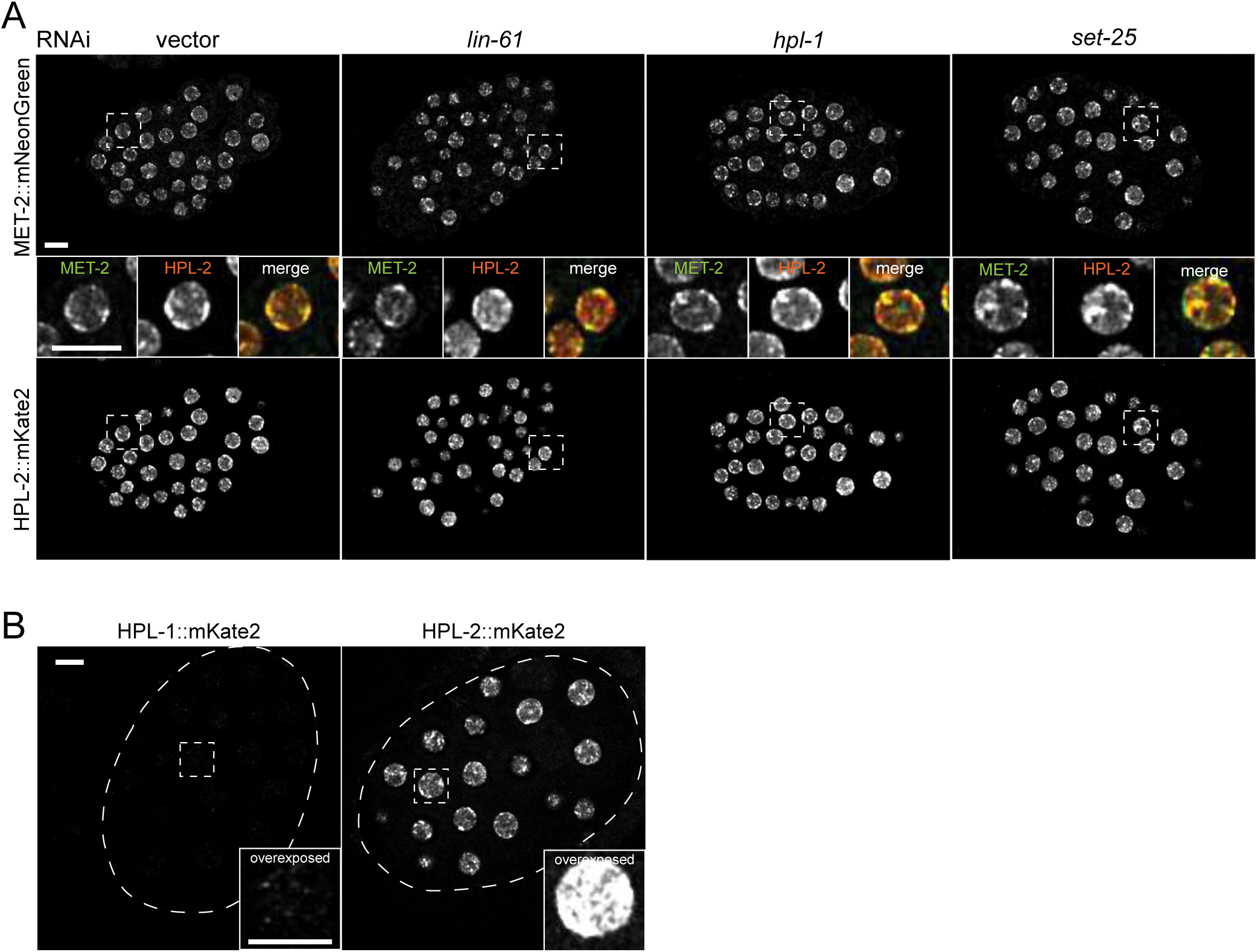
MET-2 and HPL-2 foci formation is independent of associated genes implicated in heterochromatin. (**A**) Representative images of embryos co-expressing MET-2::mNeonGreen with HPL-2::mKate2 treated with the indicated RNAi or empty vector control. Scale bar, 5 μm. N = 3, n = 30. (**B**) Representative images of embryos expressing either HPL-1::mKate2 or HPL-2::mKate2. Dashed oval indicates the circumference of the embryo. Inset shows overexposed region highlighted in the dashed box. Scale bar, 5 μm. N = 3, n = 75.

**Fig S2.**
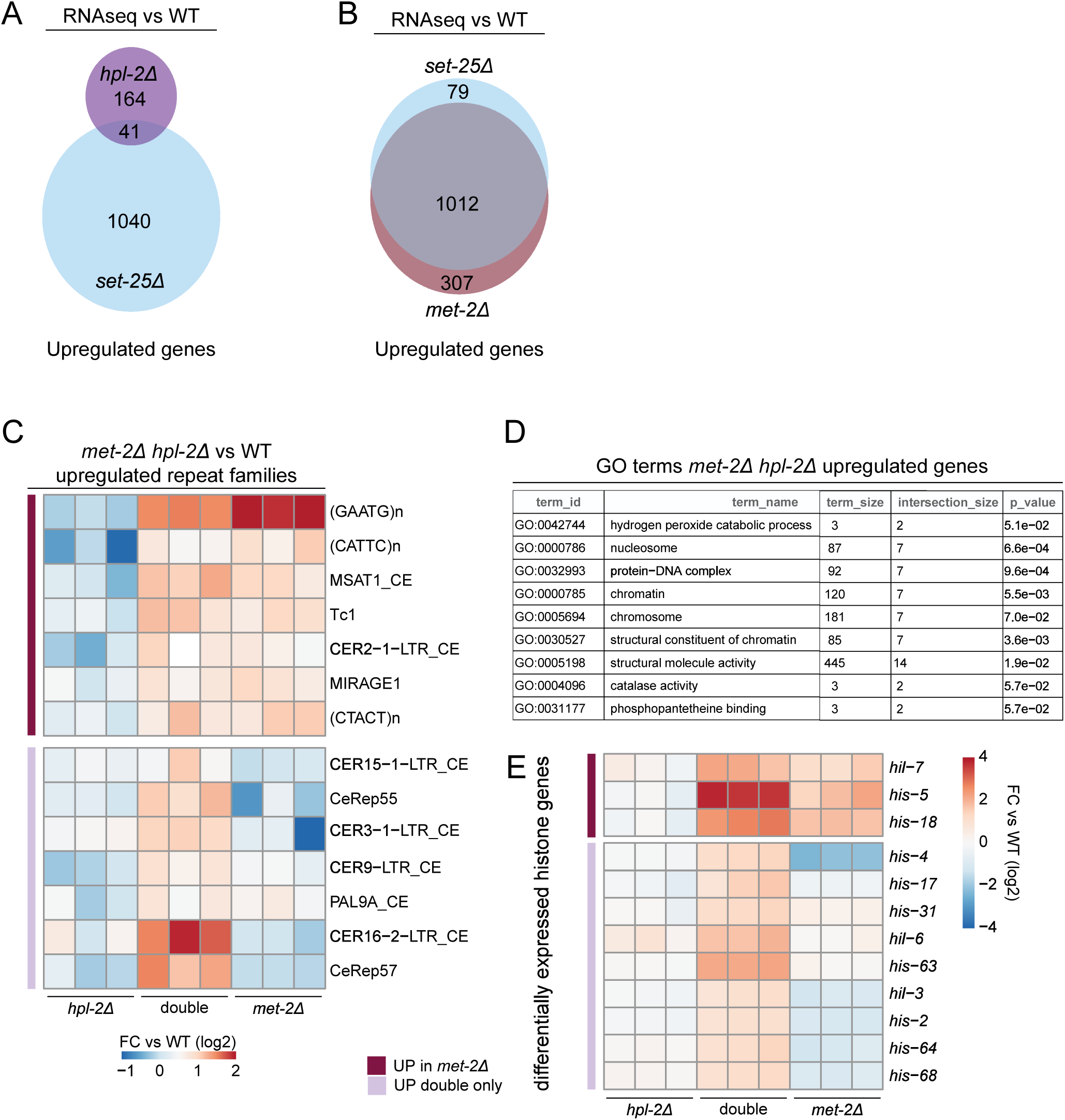
MET-2 and HPL-2 employ redundant mechanisms to silence genes and repeats. Venn diagrams showing overlap of *set-25Δ* mutant upregulated genes with (**A**) *hpl-2Δ* mutant or (**B**) *met-2Δ* mutants upregulated genes measured in independent RNAseq experiments. N = 3, FC > 2, FDR < 0.05. (**C**) Heat map showing *met-2Δ hpl-2Δ* up-regulated repeat families measured by RNAseq, and the subset of genes independently observed de-repressed in *met-2Δ* early embryos. N = 3, FC > 2, FDR < 0.05. (**D**) GO Terms from *met-2Δ hpl-2Δ* RNAseq. (**D**) Heat map showing *met-2Δ hpl-2Δ* up-regulated histone genes, as in (C).

**Fig S3.**
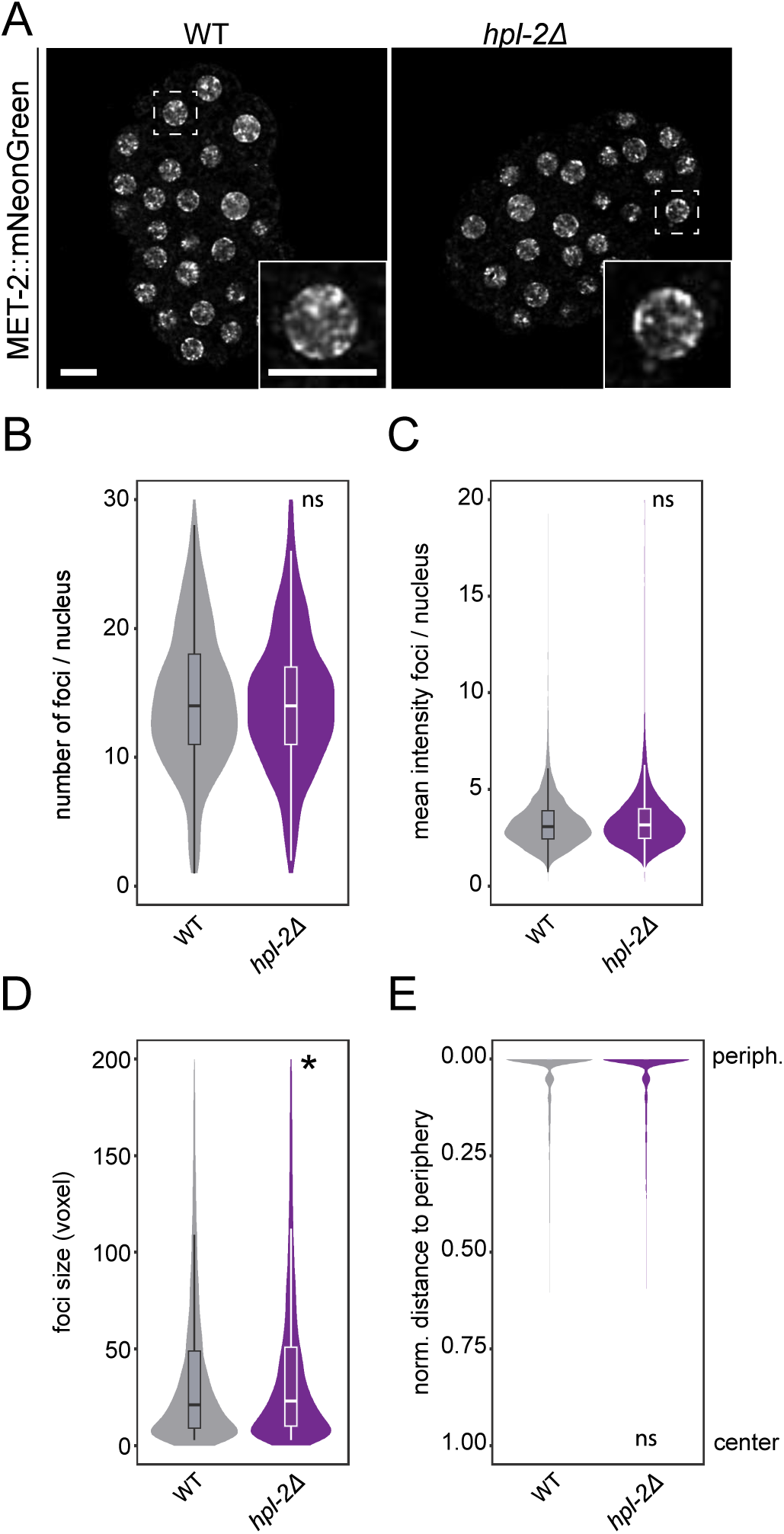
MET-2 foci are independent of *hpl-2*. (**A**) Representative live images of MET-2::mNeonGreen in WT and *hpl-2Δ* embryos. Scale bar, 5 μm. N = 3, n = 30. Quantification of MET-2::mNeonGreen foci (**B**) number, (**C**) intensity, (**D**) size, and (**E**) distance to the nuclear periphery from data in (A). *P-values (*hpl-2Δ* vs WT): (B) not significant (NS), (C) NS (D) p = 0.011, (E) NS using by one-way ANOVA, followed by Tukey post hoc test.

**Fig S4.**
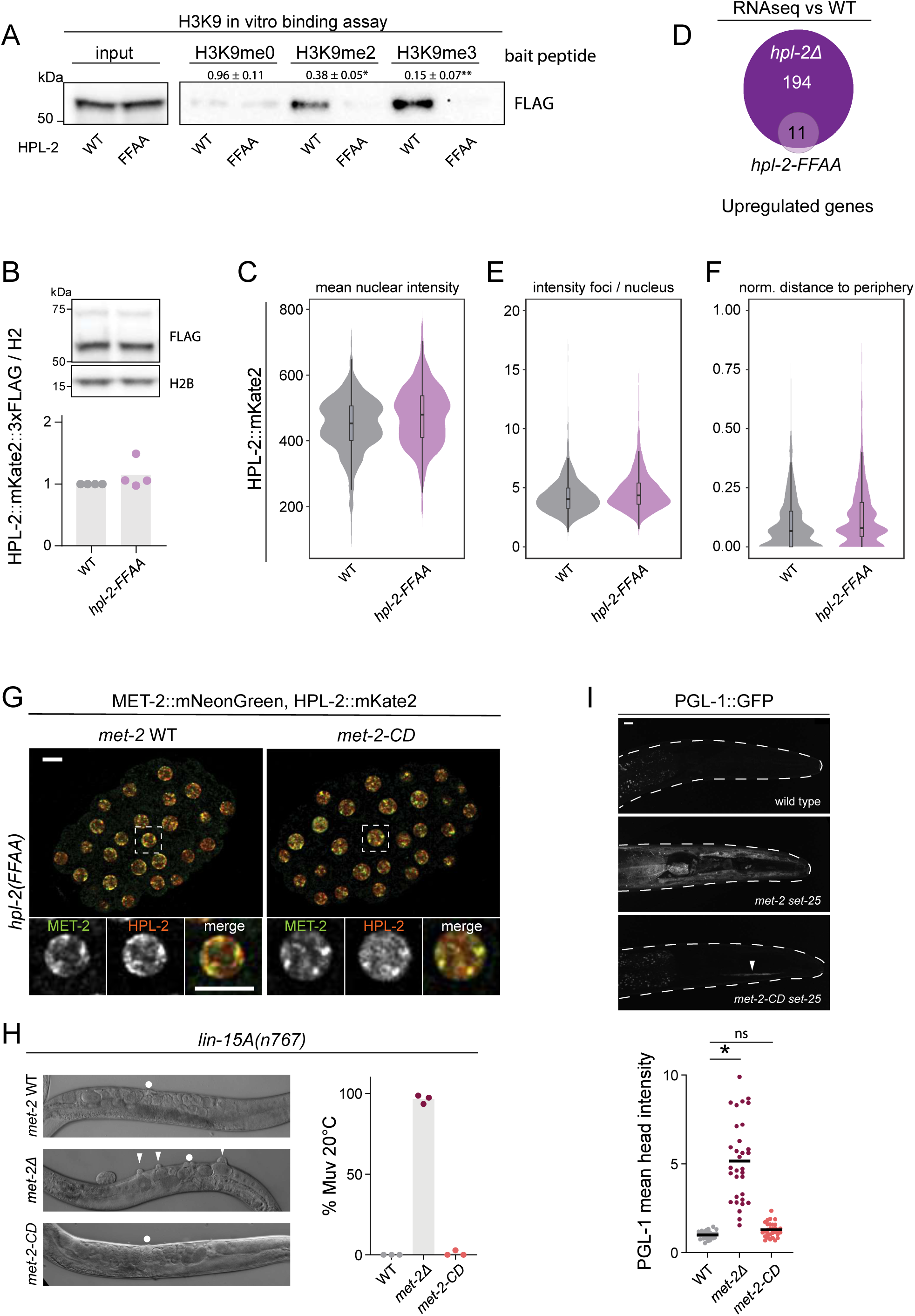
H3K9me deposition is not required for vulva specification. (**A**) Representative Western blot of H3K9me peptide in vitro binding assay. Mean +/- standard error ratio of FFAA / WT bands for each peptide is shown above the blot. N = 3. *p = 1.8 x 10^-4^, **p = 2.6 x 10^-4^ by two-sided T test. (**B**) Representative western blot of HPL-2::mKate2::3xFLAG in *hpl-2-WT* vs. *hpl-2-FFAA* protein levels. N = 4. (**C**) Image quantification in *hpl-2-WT* vs. *hpl-2-FFAA* of HPL-2::mKate2::3xFLAG nuclear intensity from Fig. 2D, p = 4.1 x 10^-3^ by one-way ANOVA, followed by Tukey post hoc test. (**D**) Venn diagram showing overlap of *hpl-2Δ* and *hpl-2-FFAA*-derepressed genes measured by RNAseq. N = 3, FC > 2, FDR < 0.05 Image quantification in *hpl-2-WT* vs. *hpl-2-FFAA* of HPL-2::mKate2::3xFLAG (**E**) foci intensity, and (**F**) distance to the nuclear periphery from Fig. 3D. P-values (*hpl-2-FFAA* vs *hpl-2-WT*): (E) not significant, (F) p = 0.039 by one-way ANOVA, followed by Tukey post hoc test. (**G**) Representative images of MET-2::mNeonGreen co-expressed with HPL-2::mKate2 in WT, *met-2-CD*, and *hpl-2-FFAA* backgrounds. Scale bar, 5 μm. N = 3, n = 30. (**H**) Representative DIC live images and quantification of multivulva (Muv) defects in worms expressing *met-2-WT*, *met-2Δ* or *met-2-CD* in the sensitized *lin-15A(n767)* background. Normal vulvae = circle, multivulva defects = arrowheads. N = 3. n: WT = 1463, *met-2Δ* = 1141, *met-2-cd* = 1346. (**I**) Representative live image maximum projections and signal quantification of endogenously-tagged PGL-1::GFP in the heads of L4 animals. Body is outlined in dashed line; mouth is to the right. Arrowhead = faint ectopic expression. Scale bar = 10 μm, N = 3, n = 30. *p < 10^-5^, ns = not significant by one-way ANOVA, followed by Tukey post hoc test.

**Fig. S5.**
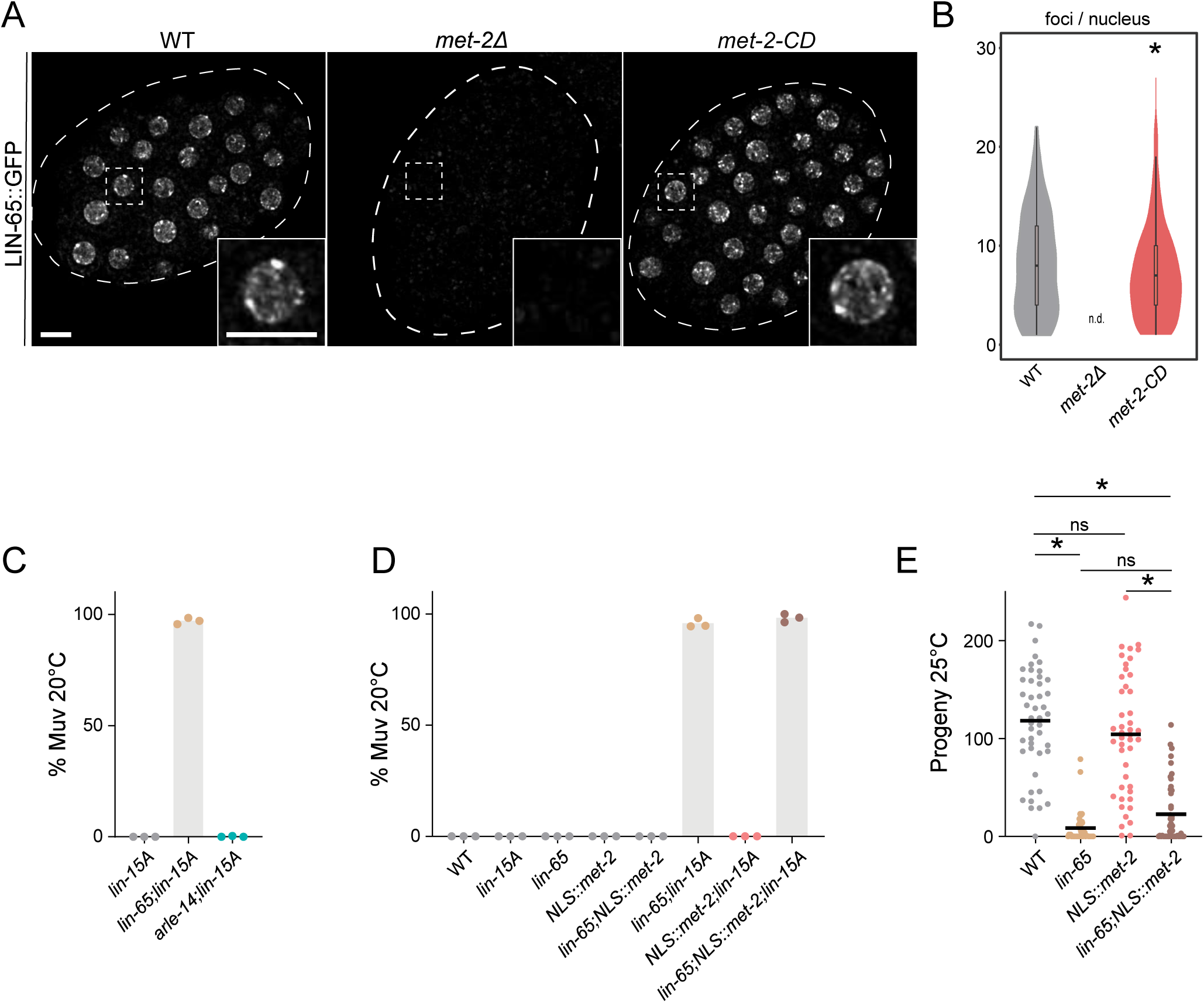
Interaction with LIN-65 is essential for MET-2 function specifying the vulva. (A) Representative live images of endogenously tagged LIN-65::GFP in WT, *met-2Δ*, and *met-2-CD* embryos. Scale bar, 5 μm. N = 3, n = 45. Dashed lines indicate embryo circumference. (**B**) Quantification of foci number from (A). n.d. = not detected, met-2-CD vs WT p = 0.028 using two-sided Wilcoxon signed-rank test. (**C**) Quantification of multivulva (Muv) defects in *lin-65Δ* or *arle-14Δ* mutants in the sensitized *lin-15A(n767)* background. N = 3. n: *lin-15A* = 1159, *lin-65Δ;lin-15A* = 1199, *arle-14Δ;lin-15A* = 903. (**D**) Quantification of multivulva (Muv) defects in worms expressing an endogenously tagged nuclear localized NLS::met-2 in the presence or absence of *lin-65* and *lin-15A*. N = 3. n: WT = 429, *lin-15A* = 404, *lin-65Δ* = 434, *NLS::met-2* = 548, *lin-65Δ;NLS::met-2* = 467, *lin-65Δ;lin-15A* = 650, *NLS::met-2;lin-15A* = 589, *lin-65Δ;NLS::met-2;lin-15A* = 692 . (**E**) Progeny number per worm 25°C in WT, *lin-65Δ*, NLS::met-2, and *lin-65Δ;NLS::met-2*. *N* = 3, n = 45 egglayers. *p < 10^-5^, ns = not significant by by one-way ANOVA, followed by Tukey post hoc test.

## Notes

https://www.ncbi.nlm.nih.gov/geo/query/acc.cgi

### Competing Interest Statement

The authors have declared no competing interest.

### Summary of Updates

This version of the manuscript adds one more author to one of the grants, which is critical for funding purposes: Jan Padeken & Lukas Stelzl are supported by the SFB 1551 Project No. 464588647 of the DFG (Deutsche Forschungsgemeinschaft).

